# The digestive system is a potential route of 2019-nCov infection: a bioinformatics analysis based on single-cell transcriptomes

**DOI:** 10.1101/2020.01.30.927806

**Authors:** Hao Zhang, Zijian Kang, Haiyi Gong, Da Xu, Jing Wang, Zifu Li, Xingang Cui, Jianru Xiao, Tong Meng, Wang Zhou, Jianmin Liu, Huji Xu

**Affiliations:** Department of Rheumatology and Immunology, Changzheng Hospital, Second Military Medical University, 200003 Shanghai, China; Peking-Tsinghua Center for Life Sciences, TsinghuaUniversity, Beijing, P.R. China; Beijing Tsinghua Changgeng Hospital, School of Clinical Medicine, Tsinghua University, 100084 Beijing, China; Department of Orthopaedic Oncology, Changzheng Hospital, Second Military Medical University, 200003 Shanghai, China; Department of Neurosurgery, Changhai Hospital, Second Military Medical University, 200003 Shanghai, China; Depanrtment of Urology, The Third Affiliated Hospital of Second Military Medical University, 201805 Shanghai, China; Division of Spine, Department of Orthopedics, Tongji Hospital affiliated to Tongji University School of Medicine, 200065 Shanghai, China; Tongji University Cancer Center, School of Medicine, Tongji University, 200092 Shanghai, China; Qiu-Jiang Bioinformatics Institute, 200003 Shanghai, China

## Abstract

Since December 2019, a newly identified coronavirus (2019 novel coronavirus, 2019-nCov) is causing outbreak of pneumonia in one of largest cities, Wuhan, in Hubei province of China and has draw significant public health attention. The same as severe acute respiratory syndrome coronavirus (SARS-CoV), 2019-nCov enters into host cells via cell receptor angiotensin converting enzyme II (ACE2). In order to dissect the ACE2-expressing cell composition and proportion and explore a potential route of the 2019-nCov infection in digestive system infection, 4 datasets with single-cell transcriptomes of lung, esophagus, gastric, ileum and colon were analyzed. The data showed that ACE2 was not only highly expressed in the lung AT2 cells, esophagus upper and stratified epithelial cells but also in absorptive enterocytes from ileum and colon. These results indicated along with respiratory systems, digestive system is a potential routes for 2019-nCov infection. In conclusion, this study has provided the bioinformatics evidence of the potential route for infection of 2019-nCov in digestive system along with respiratory tract and may have significant impact for our healthy policy setting regards to prevention of 2019-nCoV infection.

## Introduction

At the end of 2019, a rising number of pneumonia patients with unknown pathogen has been emerging in one of largest cities of China, Wuhan, and quickly spread throughout whole country[1]. A novel coronavirus was then isolated from the human airway epithelial cells and was named 2019 novel coronavirus (2019-nCoV)[2]. The complete genome sequences has reveled that 2019-nCoV sharing 86.9% nucleotide sequence identity to a severe acute respiratory syndrome (SARS)-like coronavirus detected in bats (bat-SL-CoVZC45, MG772933.1). This suggested that 2019-nCoV is the species of SARS related coronaviruses (SARSr-CoV) by pairwise protein sequence analysis[2, 3].

As for the clinical manifestations of 2019-nCoV infection, fever and cough are most common symptoms at onset[4, 5]. In addition, it frequently induces severe enteric symptoms, such as diarrhea and nausea, which are even graver than those of SARS-CoV and Middle East respiratory syndrome coronavirus (MERS-CoV)[6, 7]. However, a little was known why and how the 2019-nCov induced enteric symptoms. In addition, it is unknown yet whether 2019-nCoV can be transmitted through the digestive tract besides respiratory tract[5].

The prerequisite of coronaviruses infection is its entrance into the host cell. During this process, the spike (S) glycoprotein recognizes host cell receptors and induces the fusion of viral and cellular membranes[8]. In 2019-nCoV infection, a metallopeptidase, angiotensin converting enzyme II (ACE2) is proved to be the cell receptor, the same as SARS-CoV infection[9-11]. 2019-nCoV can enter into ACE2-expressing cells, but not into cells without ACE2 or cells with other coronavirus receptors, such as aminopeptidase N and dipeptidyl peptidase[10]. Thus, ACE2 plays an vital role in the 2019-nCoV infection.

In order to explore the infection routes of 2019-nCov and the roles of ACE2 in digestive system infection, we identified the ACE2-expressing cell composition and proportion in normal human lung and gastrointestinal system by single-cell transcriptomes based on the public databases. A striking finding is that ACE2 was not only expressed in lung AT2 cells, but also found in esophagus upper and stratified epithelial cells and absorptive enterocytes from ileum and colon. In addition, the enteric symptoms of 2019-nCov may be associated with the invaded ACE2-expressing enterocytes. These findings indicate that the digestive systems along with respiratory tract may be potential routes of 2019-nCov infection may have significant impact for our healthy policy setting regards to prevention of 2019-nCoV infection..

## Materials and Methods

### Data Sources

Single-cell expression matrices for the lung, esophagus, stomach, ileum and colon were obtained from the Gene Expression Omnibus (GEO; https://www.ncbi.nlm.nih.gov/)[12], Single Cell Portal (https://singlecell.broadinstitute.org/single_cell) and Human Cell Atlas Data Protal. (https://data.humancellatlas.org). Single-cell data for the esophagus and lung were obtained from the research by E Madissoon et al which contained 6 esophageal and 5 Lung tissue samples[13], which contained 6 esophageal and 5 lung tissue samples.. The data of gastric mucosal samples from 3 non-atrophic gastritis and 3 chronic atrophic gastritis patients were obtained from GSE134520[14]. GSE134809[15] was comprised of 22 ileal specimens from 11 ileal Crohn’s disease patients and only non-inflammatory samples were selected for analysis. The research by Christopher S et al[16] included 12 normal colon samples.

### Quality Control

Low quality Cells with expressed genes were lower than 200 or larger than 5000 were removed. We further required the percentage of UMIs mapped to mitochondrial or ribosomal genes to be lower than 20%.

### Data Integration, Dimension Reduction and Cell Clustering

Different data processing methods were performed for different single-cell projects according to the downloaded data.

#### Esophagus and lung datasets

Seurat [17] rds data was directly download from supplementary material in the research by E. Madissoon al [13]. Uniform Manifold Approximation and Projection (UMAP) visualization were performed for gaining clusters of cells.

#### Stomach and ileum datasets

Single cell data expression matrix was processed with the R package Seurat (version 3.0)[17]. We first utilized ‘‘NormalizeData’’ normalize and the single-cell gene expression data. UMI counts were normalized by the total number of UMIs per cell, multiplied 10000 for the normalization and were transformed to the log-transformed counts. The highly variable Genes (HVGs) were identified using the function “FindVariableGenes”. We then used “FindIntegrationAnchors” and “Integratedata” function to merge multiple sample data within each dataset. After removing unwanted sources of variation from a single-cell dataset such as cell cycle stage, or mitochondrial contamination, we used the ‘‘RunPCA’’ function to perform the principle component analysis (PCA) on the single-cell expression matrix with significant HVGs. Then we constructed a K-nearest-neighbor graph based on the euclidean distance in PCA space using the “FindNeighbors” function and applied Louvain algorithm to iteratively group cells together by “FindClusters” function with optimal resolution. UMAP was used for visualization purposes.

#### Colon Dataset

Single cell data expression matrix was processed with the R package LIGER[18] and Seurat[17]. We first normalized the data to account for differences in sequencing depth and capture efficiency among cells. Then we used “selectGenes” function to identify variable genes on each dataset separately and took the union of the result. Next integrative non-negative matrix factorization was performed to identify shared and distinct metagenes across the datasets and the corresponding factor loadings for each cell using “optimizeALS” function in LIGER. We selected a k of 15 and lambda of 5.0 get a plot of expected alignment. We then identified clusters shared across datasets and aligned quantiles within each cluster and factor using “quantileAlignSNF” function. Next nonlinear dimensionality reduction was performed using “RunUMAP” function in Seurat and the results were visualized with UMAP.

### Identification of cell types and Gene expression analysis

We annotated cell clusters based on the expression of known cell marker and the clustering information provided in the articles. Then we used “RunALRA” function in Seurat to imput dropped out values in scRNA-seq data. Feature plots and violin plots were generated using Seurat to show imputed gene expression. In order to compare gene expression in different datasets, we used “Quantile normalization” in R package preprocessCore (R package version 1.46.0. https://github.com/bmbolstad/preprocessCore) to preprocess data. Then gene expression data were further denoised by adding random generation for the normal distribution with mean equal to mean and standard deviation equal to sd.

## Results

### Annotation of cell types

The gastrointestinal system is composed of esophagus, stomach, ileum, colon and cecum. In this study, 4 datasets with single-cell transcriptomes of esophagus, gastric, ileum and colon were analyzed, along with lung (Additional file). Based on Cell Ranger output, the gene expression count matrices were used to present sequential clustering of cells according to different organs or particular clusters. The cell type identity in each cluster was annotated by the expression of the known cell type markers.

In the esophagus, 14 cell types were identified through 87,947 cells. Over 90% cells fall into four major epithelial cell types: upper, stratified, suprabasal, and dividing cells of the suprabasal layer (Fig. 1A). The additional cells from the basal layer of epithelia clustered more closely to the gland duct and mucous secreting cells. Lymph vessel and endothelial cells are associated with vessel tissues. Immune cells in the esophagus include T cells, B cells, monocytes, macrophages, dendritic cells (DCs), and mast cells.

**Figure 1:**
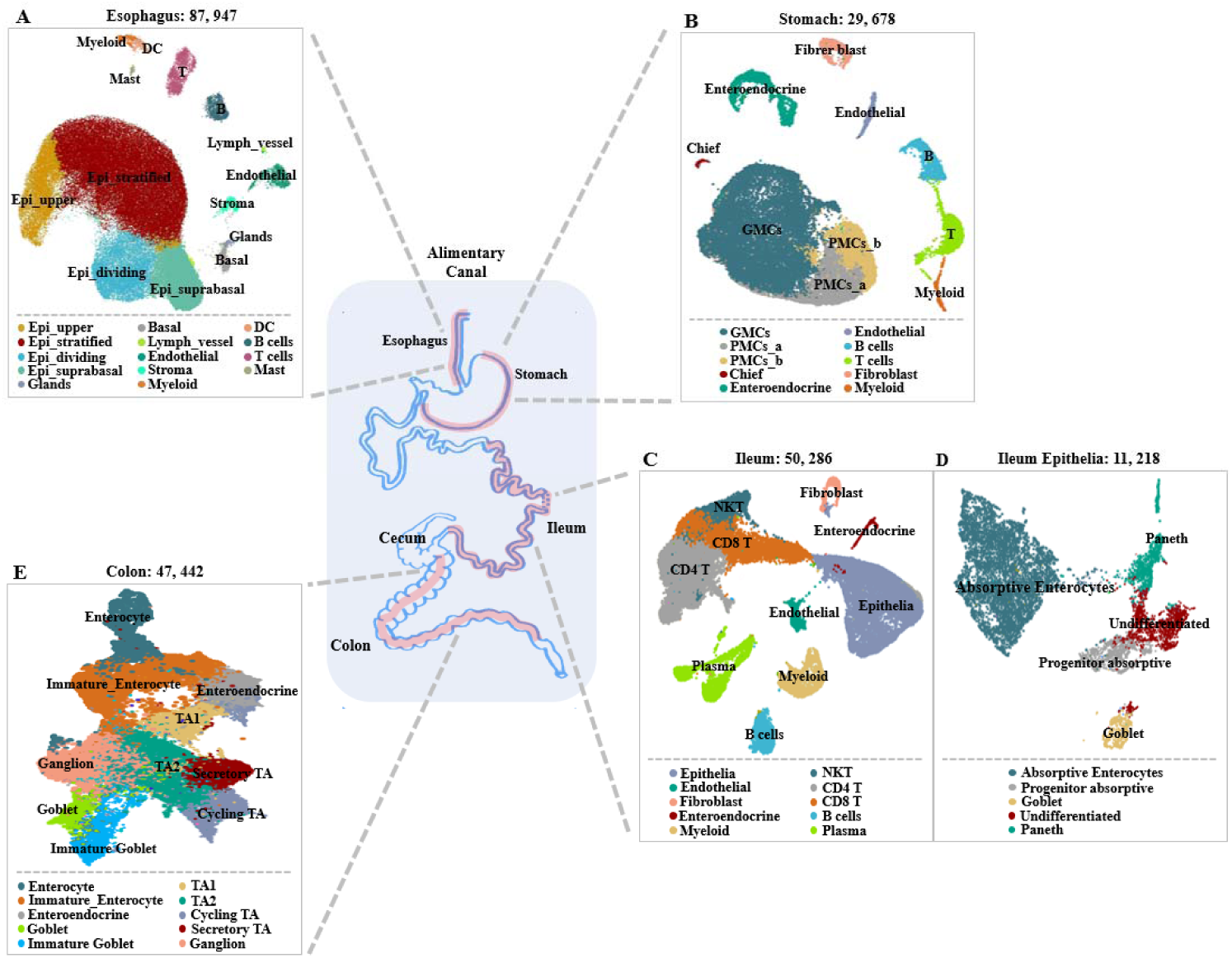
Single-Cell Atlas of digestive tract samples. (A). The UMAP plot of 87947 esophageal cells to visualize cell-type clusters (B). The UMAP plot of 29678 gastric mucosa cells to visualize cell-type clusters. (C). The UMAP plot of 50286 ileal cell cells to visualize cell-type clusters. (D). The UMAP plot of 11218 ileal epithelial cells to visualize finer clusters. Epithelial cells in ileum were further divided into finer cell subsets because of the heterogeneity within the cell population according to transcription characteristics. (E). The UMAP plot of 47442 colon cells to visualize cell-type clusters.

A total of 29,678 cells and 10 cell types were identified in the stomach after quality control with a high proportion of gastric epithelial cells, including antral basal gland mucous cells (GMCs), pit mucous cells (PMCs), chief cells and enteroendocrine cells (Fig. 1B). The non-epithelial cell lineages were composed of T cells, B cells, myeloid cells, fibroblasts and endothelial cells.

After quality controls, 50,286 cells and 10 cell types were identified in the ileum (Fig. 1C). The detected cell types included epithelia, endothelial, fibroblast and enteroendocrine cells. The identified immune cell types were myeloid, CD4^+^T, CD8^+^T and natural killer T (NKT) cells, along with plasma and B cells. Among 11,218 epithelial cells, 5 cell types were identified, namely, absorptive enterocytes, progenitor absorptive, goblet, Paneth and undifferentiated cells (Fig. 1D).

All the 47,442 cells from the colon were annotated after quality controls (Fig. 1E). Absorptive and secretory clusters were identified in epithelial cells. The absorptive clusters included further sub-clusters for transit amplifying (TA) cells (TA 1, TA 2), immature enterocytes, and enterocytes. The secretory clusters included sub-clusters for progenitor cells (secretory TA, immature goblet) and for mature cells (goblet, and enteroendocrine). Ganglion cells and cycling TA cells were also identified in the final UMAP.

### Cell type-specific ACE2 expression

With regard to stomach, the expression of ACE2 is relatively low in all the clusters (Fig. 2B, C). The selected cell type-specific marker genes were used to identify each cluster in the stomach (Fig. 2C). MUC6 and TIFF1 were highly expressed in all the clusters. PGA4 was used to identify chief cells, along with CHGB for enteroendocrine cells, CD34 for endothelial cells, CD79A for B cells, CD8A and PRF1 for T cells, VCAN and COL1A1 for fibrosus blast and CLEC10A for myeloid cells.

**Figure 2.**
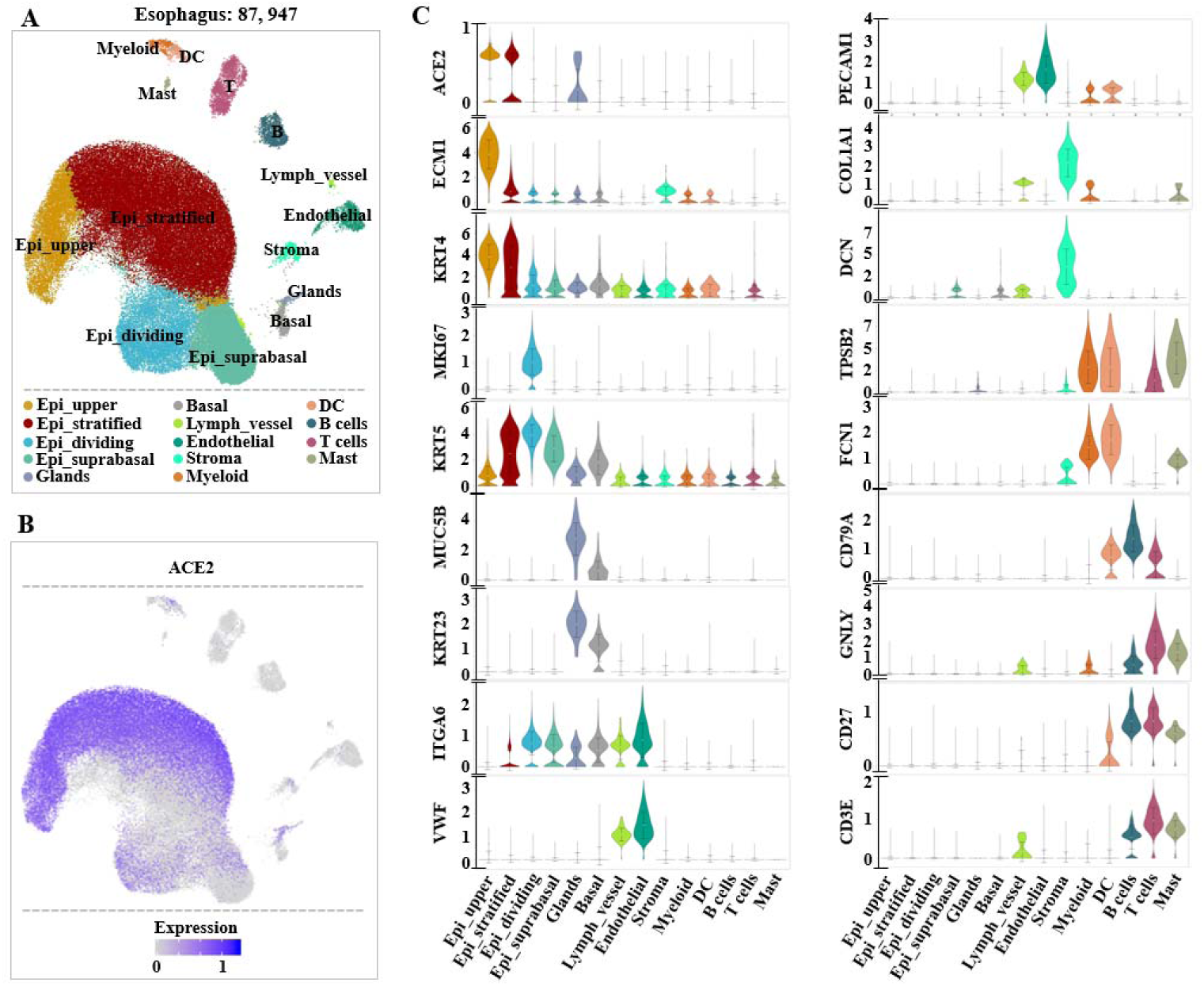
Single-cell analysis of esophageal cells. (A). UMAP plots showing the landscape of esophageal cells. 14 cell clusters were identified across 87947 cells. (B). UMAP plots showing the expression of *ACE2* across clusters. (C).Violin plots for esophageal clusters marker genes and *ACE2* across clusters. The expression is measured as the log_2_ (TP10K+1).

As for esophagus, ACE2 was highly expressed in upper and stratified epithelial cells (Fig. 3B, C). The glands also have a low expression of ACE2 (Fig. 3C). The selected cell type-specific marker genes were used to identify each cluster in the esophagus (Fig. 3C). ECM1 was highly expressed in upper epithelial cells. KRT4 and 5 were mainly found in stratified epithelial cells. KI67 was used to identify dividing epithelial cells, with MUC5B and KRT23 for glands, COL1A1 and DCN for stroma cells, VWF and PECAM1 for lymph vessel and endothelial cells. TPSB2, FCN1, CD79A, GNLY, CD27 and CD3E were used for immune cells, such as myeloid, DC, B, T and mast cells.

**Figure 3.**
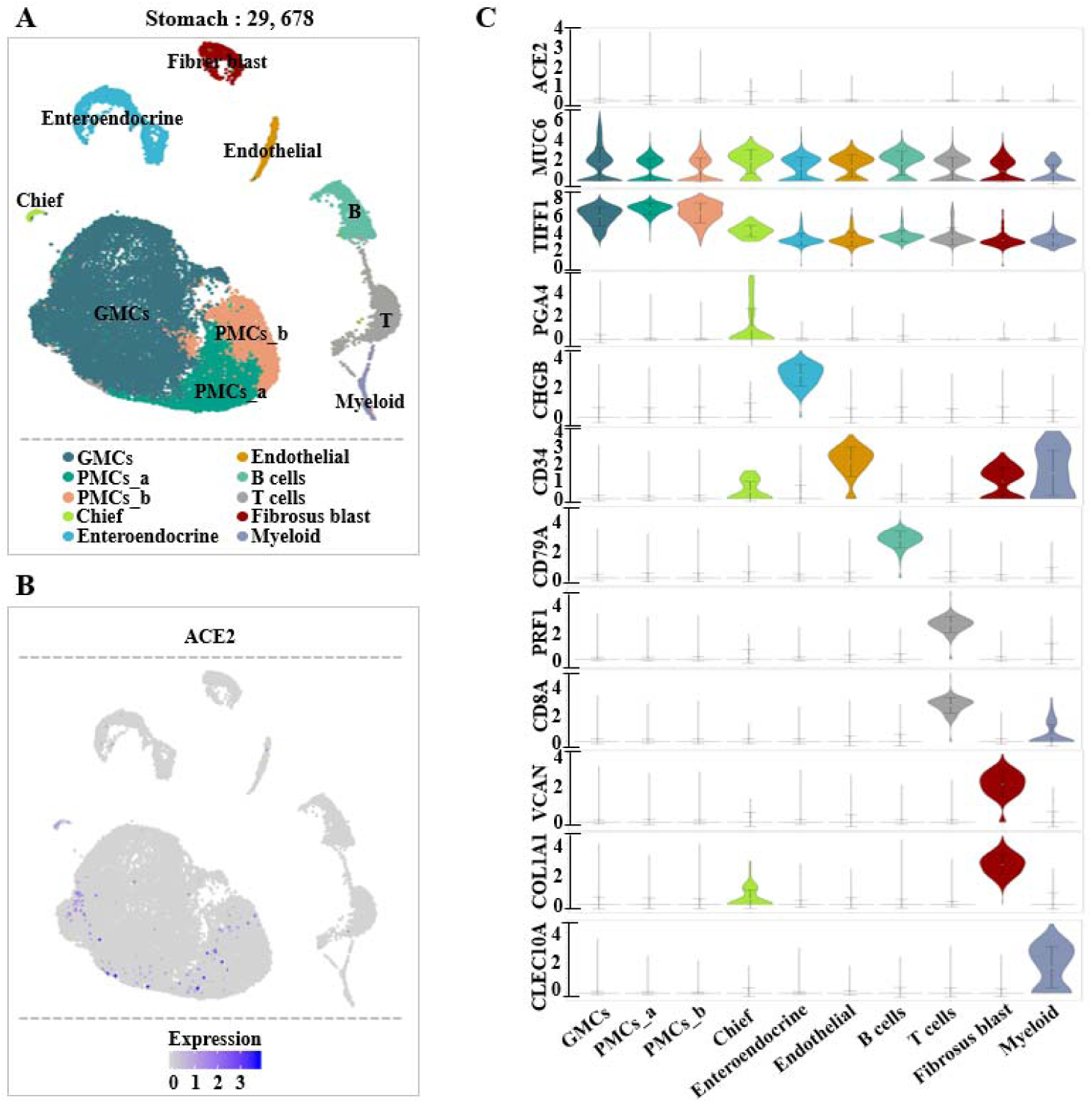
Single-cell analysis of gastric mucosal cells. (A). UMAP plots showing the landscape of gastric mucosal tissue. 10 cell clusters were identified across 29678 cells after quality control, dimensionality reduction and clustering. (B). UMAP plots showing the expression (grey to blue) of gene *ACE2* across clusters. (C).Violin plots for gastric mucosal clusters marker genes and *ACE2* across clusters. The expression is measured as the log_2_ (TP10K+1).

In the epithelial cells of the ileum, ACE2 was highly expressed in absorptive enterocytes and less expressed in progenitor absorptive cells, which was similar to those in the colon (Fig. 4B, C). The selected cell type-specific marker genes were also used to identify the epithelial cells of the ileum (Fig. 4C). SEC2A5 was found mainly in the absorptive enterocytes and progenitor absorptive cells. CD24 was found in all epithelial cells except absorptive enterocytes. MII67 and AMACR were highly expressed in undifferentiated and Paneth cells, respectively. BCAS1 was used to identify undifferentiated cells, with AMACR for Paneth.

**Figure 4.**
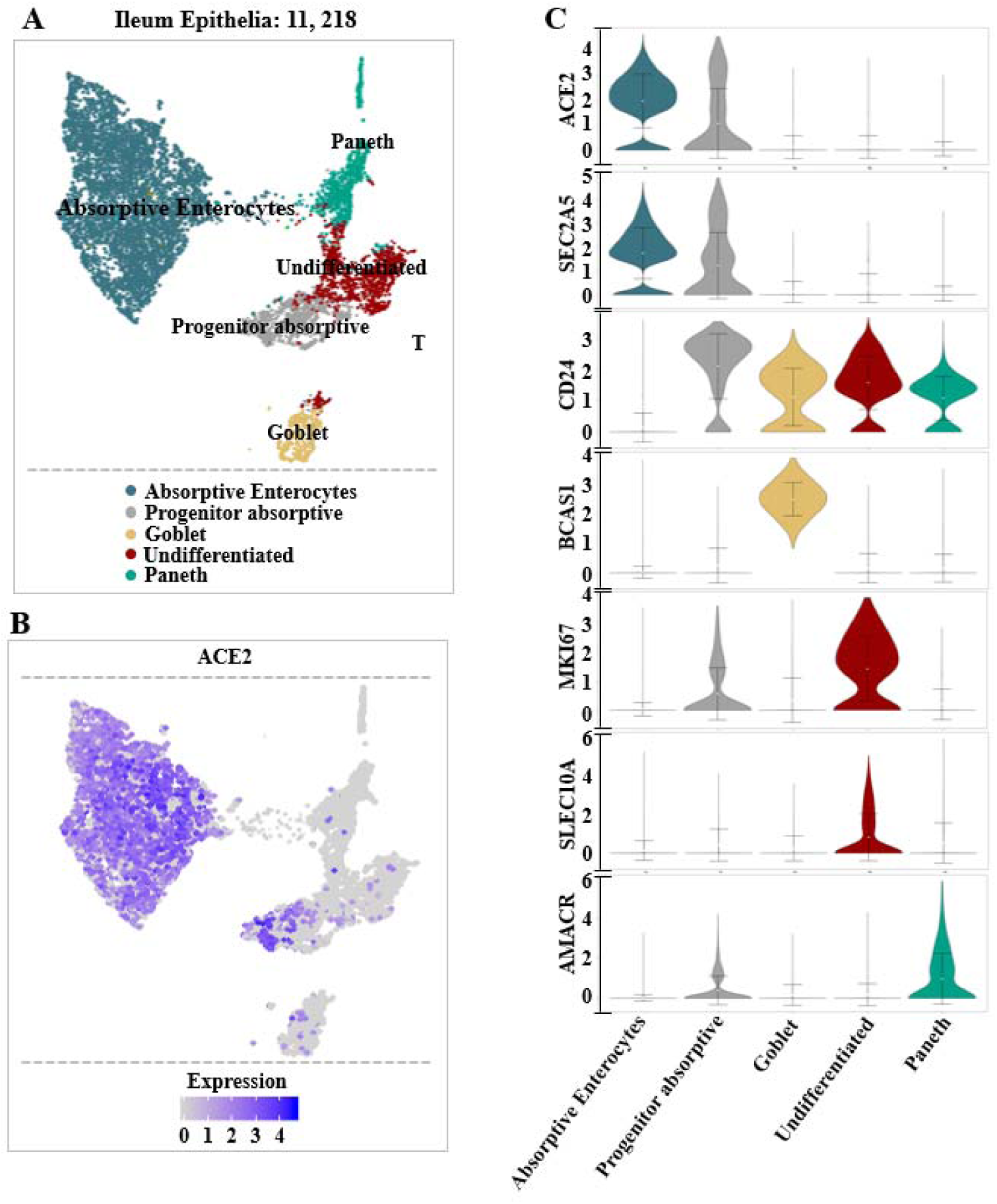
Single-cell analysis of ileal epithelial cells. (A). UMAP plots showing the landscape of ileal epithelial cells. 10 cell clusters were identified across 11218 cells after quality control, dimensionality reduction and clustering. (B). UMAP plots showing the expression of *ACE2* across clusters. (C). Violin plots for ileal epithelial marker genes and *ACE2* across clusters. The expression is measured as the log_2_ (TP10K+1).

In the colon, ACE2 was mainly found in enterocytes and less expressed in immature enterocytes (Fig. 5B, C). The selected cell type-specific marker genes were used to identify each cluster in the colon (Fig. 5C). AQPB was mainly found in enterocytes and immature enterocytes. Additionally, ZG16 and ITLN1 was highly expressed in goblet and immature goblet. The expression of APOE was in TA2 and secretory TA. CD27 and TPH1 were used to identify enteroendocrine, with SPC25 for cycling TA. After initial quality controls, 57,020 cells and 25 cell types were identified in the lung (Fig. 6A). The detected cell types included ciliated, alveolar type 1 (AT1) and alveolar type 2 (AT2) cells, along with fibroblast, muscle, and endothelial cells. The identified immune cell types were T, B and NK cells, along with macrophages, monocytes and dendritic cells (DC). ACE2 was mainly expressed in AT2 cells and could also be found in AT1 and fibroblast cells (Fig. 6B).

**Figure 5.**
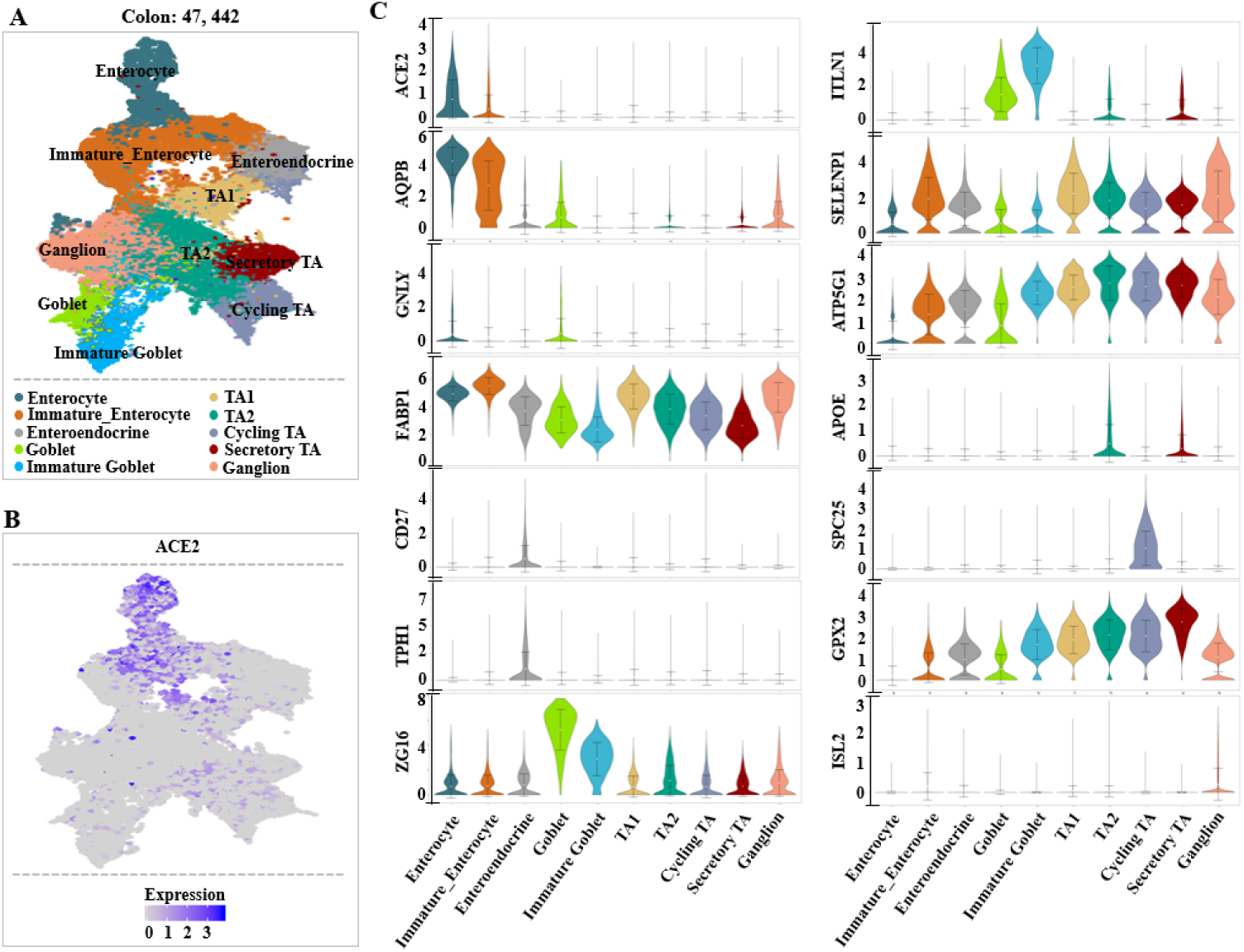
Single-cell analysis of colon cells. (A). UMAP plots showing the landscape of colon cell cells. 10 cell clusters were identified across 47442 cells after quality control, dimensionality reduction and clustering. (B). UMAP plots showing the expression of *ACE2* across clusters. (C). Violin plots for colon clusters marker genes and *ACE2* across clusters. The expression is measured as the log_2_ (TP10K+1).

**Figure 6.**
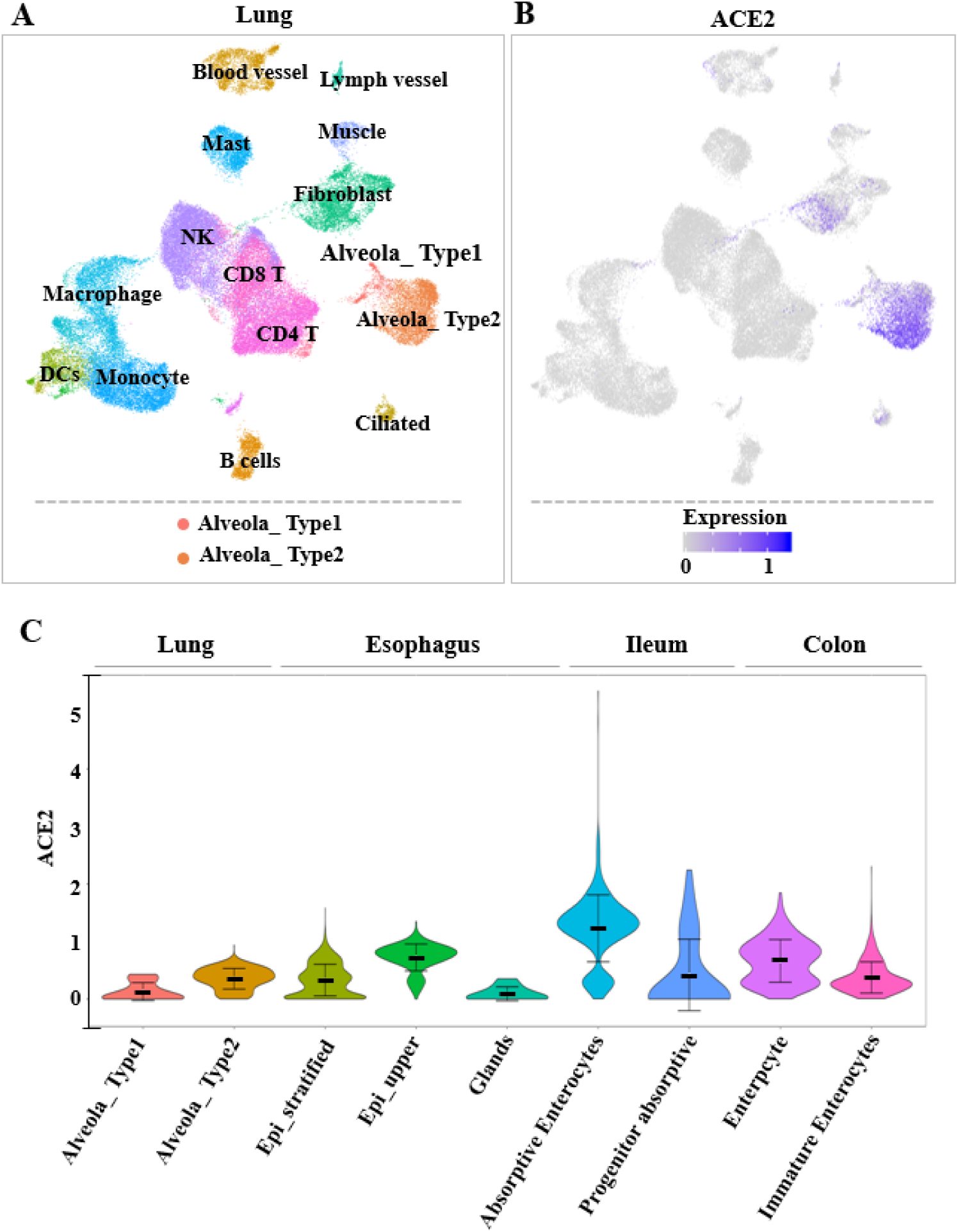
Single-cell analysis of lung cells. (A). UMAP plots showing the landscape of lung cells. 16 cell clusters were identified across 57020 cells. (B). UMAP plots showing the expression of *ACE2* across lung clusters. (C). Violin plots for *ACE2* across 2 lung clusters and 7 digestive tract clusters. Gene expression matrix was normalized and denoised to remove unwanted technical variability across 4 datasets.

Among all the ACE2-expressing cells in normal digestive system and lung, the expression of ACE2 was more in ileum and colon than that in the lung and esophagus (Fig. 6C).

## Discussion

The coronaviruses is the common infection source of upper respiratory, gastrointestinal and central nervous system in humans and other mammals[19]. At the beginning of the twenty-first century, two betacoronaviruses, SARS-CoV and MERS-CoV, caused persistent public panics and became the most significant public health events[20]. In December 2019, a novel identified coronavirus (2019-nCov) induced an ongoing outbreak of pneumonia in Wuhan, Hubei, China with arising number of infected patients[4]. Till now, its infection routes and digestive system infection are still unclear. In this study, we found the high expressions of ACE2, the cell entry receptor of 2019-nCov, in the lung AT2 cells, esophagus upper and stratified epithelial cells and absorptive enterocytes from ileum and colon, indicating that not only respiratory system but also digestive system are potential routes of infection. In addition, the enteric symptoms of 2019-nCov may be associated with the invaded ACE2-expressing enterocytes.

Generally, many respiratory pathogens, such as influenza, SARS-CoV and SARSr-CoV, cause enteric symptoms, so is 2019-nCov[4, 5]. As a classic respiratory coronavirus, SARS often causes enteric symptoms along with respiratory symptoms. Moreover, transmission with stool is also a neglected risk for SARS[21]. During the infection of SARS and highly pathogenic strains of influenza, their enteric symptoms are associated with the increased permeability to intestinal lipopolysaccharide (LPS) and bacterial transmigration through gastrointestinal wall[22, 23]. However, the mechanism of 2019-nCov-induced enteric symptom is still unknown.

A recent study revealed that similar to SARS-CoV and MERS-CoV, ACE2 was the cell entry receptor for 2019-nCov[10]. Previously, ACE2 was isolated from SARS-CoV-permissive Vero E6 cells[24]. It could interact with a defined receptor-binding domain (RBD) of CTD1 in SARS-CoV and facilitate efficient cross-species infection and person-to-person transmission[9, 25]. The “up” and “down” transition of CTD1 allows ACE2 binding by regulating the relationship among CTD1, CTD2, S1-ACE2 complex and S2 subunit[26]. With regard to human HeLa cells, expressing ACE2 from human, civet, and Chinese horseshoe bat can help many kinds of SARSr-CoV, including 2019-nCov, to enter into the cells, indicating the important role of ACE2 in cellular entry [10, 27-29]. Therapeutically, anti-ACE2 antibody can block viral replication on Vero E6 cells[24].

By analyzing the expression of ACE2 in normal human gastrointestinal system and lung, we found high expression of ACE2 in the lung AT2 cells, esophagus upper and stratified epithelial cells and absorptive enterocytes from ileum and colon. Similar to the previous study, ACE2 was more expressed in AT2 cells and less expressed in AT1 cells in normal lung[30]. In lung alveoli, AT1 epithelial cells are responsible for gas exchange and AT2 cells are in charge of surfactant biosynthesis and self-renewing[31]. In SARS-CoV infection, AT2 is the major infected cell types by viral antigens and secretory vesicles detection. Its expression in AT2 cells is variable in different donors, which may be associated with different susceptibility and seriousness[30]. Thus, we suppose that AT2 cells might be the key 2019-nCov-invaded cell in lung and its number might be associated with the severity of respiratory symptoms, which can explain the existence of asymptomatic 2019-nCov carrier.

ACE2 was also highly expressed in the esophagus upper and stratified epithelial cells. Histologically, both esophagus and respiratory system organs, such as trachea and lung are originated from the anterior portion of the intermediate foregut[32]. After being separated from the neighboring respiratory system, the esophagus undergoes subsequent morphogenesis of a simple columnar-to-stratified squamous epithelium conversion[33]. The stratified squamous epithelium can be nourished by submucosal glands and sustain the passing of the abrasive raw food. In Barrett’s oesophagus (BE), acid reflux-induced oesophagitis and the multilayered epithelium (MLE) are associated with both upper and stratified epithelial cells[34].

In the digestive system, besides esophagus upper and stratified epithelial cells, ACE2 was also found in the absorptive enterocytes from ileum and colon, the most vulnerable intestinal epithelial cells. In microbe infections, the intestinal epithelial cells function as a barrier and help to coordinate immune responses[35]. The absorptive enterocytes can be infected by coronavirus, rotavirus and noroviruses, resulting in diarrhea by destructing absorptive enterocytes, malabsorption, unbalanced intestinal secretion and activated enteric nervous system[36-38]. Thus, we suppose that the enteric symptom of diarrhea might be associated with the invaded ACE2-expressing enterocytes. In addition, due to the high expression of cell receptor ACE2 in esophagus upper and stratified epithelial cells and absorptive enterocytes from ileum and colon, we suppose that digestive system can be invaded by 2019-nCov and serve as a route of infection.

## Conclusion

This study provides the bioinformatics evidence for the potential respiratory and digestive systems infection of 2019-nCov and assists clinicians in preventing and treating the 2019-nCoV infection.

